# Characterization of clinical significance of PD-1/PD-Ls expression and methylation in patients with low grade glioma

**DOI:** 10.1101/2020.03.20.999573

**Authors:** Jie Mei, Yun Cai, Rui Xu, Xuejing Yang, Weijian Zhou, Huiyu Wang, Chaoying Liu

## Abstract

**Background:** Immune checkpoints play crucial roles in immune escape of cancer cells. However, the exact prognostic values of expression and methylation of programmed death 1 (PD-1), programmed death-ligand 1 (PD-L1) and PD-L2 in low-grade glioma (LGG) have not been defined yet.

**Methods:** A total of 514 LGG samples from TCGA dataset containing both PD-1, PD-L1 and PD-L2 expression, DNA methylation, and survival data were enrolled into our study. The clinical significance of PD-1/PD-Ls expression and methylation in LGG were explored. Besides, the correlation between PD-1/PD-Ls expression and methylation with the infiltration levels of tumor-infiltrating immune cells (TIICs) was assessed. Moreover, GO enticement analysis of PD-1/PD-Ls co-expressed genes was performed as well. R 3.6.2 and GraphPad Prism 8 were applied as main tools for the statistical analysis and graphical exhibition.

**Results:** PD-1/PD-Ls had distinct co-expression patterns in LGG tissues. The expression and methylation status of PD-1/PD-Ls seemed to be various in different LGG subtypes. Besides, upregulated PD-1/PD-Ls expression and hypo-methylation of PD-1/PD-Ls were associated with worse survival in LGG patients. In addition, PD-1/PD-Ls expression was revealed to be positively associated with TIICs infiltration, while their methylation was negatively associated with TIICs infiltration. Moreover, the PD-1/PDLs correlated gene profiles screening and Gene Ontology (GO) enrichment analysis uncovered that PD-1/PDLs and their positively correlated gene mainly participated in immune response related biological processes.

**Conclusions:** High expression and hypo-methylation of PD-1/PD-Ls significantly correlated with unfavorable survival in LGG patients, suggesting LGG patients may benefit from PD1/PD-Ls checkpoint inhibitors treatment.

## Introduction

Glioma is a common neuroepithelial-derived primary brain tumor, which is one of most fetal malignances worldwide. As a heterogeneous disease, classification of glioma is essential for therapeutic guidance and prognostic assessment, which largely relied on tumor histopathologic features [1]. World Health Organization (WHO) grade system, the most authoritative classification, divides glioma into two main classes, which contains low-grade glioma (LGG) and glioblastoma (GBM). LGG is slower growing than their high-grade counterparts, accounting for 10-20% of all primary intracranial tumors [2]. Although surgical resection is the preferred therapeutic strategy for glioma, substantial efforts has also been made to recognize the critical interplay between glioma and immunity [3, 4].

Immunotherapy is one of the most encouraging strategies for tumor treatment, and the most common therapy is to interrupt the interaction between immune checkpoints expressed on tumor and immune cells, which blocks the immune escape of tumor cells to some extent [5]. Programmed death 1 (PD-1), also termed as cluster of differentiation 279 (CD279), is an important immunosuppressive molecule expressed on T cells and other immune cells membrane, which has been widely reported across multiple malignant tumors [6]. Programmed death-ligand 1 (PD-L1) and PD-L2 are transmembrane proteins that are accounted to play critical roles in triggering the cancer immunity escape by binding to their receptor PD-1 [7, 8]. Previously, PD-L1 and PD-L2 (PD-Ls) expression have been revealed to be correlated with poor prognosis of glioma [9, 10]. As we all known, two main glioma subtypes, LGG and GBM, exhibit different biological patterns and PD-Ls expression. However, we failed to obtain an integrated study on PD-1/PD-Ls expression and methylation in LGG. Only several researches observed the promising prognostic impact of PD-L1 in GBM [11, 12]. Thus, the relationship between expression, methylation, and prognostic values of PD-1/PD-Ls in LGG need to be further explored.

In this research, to define the PD-1/PD-Ls expression and regulatory factors in LGG, we took advantage of the Cancer Genome Atlas (TCGA), including RNA-sequencing mRNA expression, DNA methylation, and copy number data. Besides, we further checked the prognostic values of PD-1/PD-Ls expression and methylation status in subpopulations receiving different therapies. This is the first integrative research that systematically characterizes PD-1/PD-Ls expression and methylation in LGG molecularly and clinically, providing a comprehensive insight into the values of PD-1/PD-Ls in predicting prognosis in patients with LGG.

## Materials and methods

### Acquisition of TCGA data

The data of RNA-sequencing (IlluminaHiSeq), DNA methylation (Methylation450k), and copy number (gistic2 thresholded) as well as clinical information in TCGA-LGG dataset were downloaded from UCSC Xena (https://xenabrowser.net/datapages/). The gene expression level was assessed as in log2(x+1) transformed RSEM normalized count. Main clinical data contained the histological type, IDH mutation status, WHO grade, therapeutic strategies, and survival information. For further analysis, a total of 514 samples containing both gene expression, DNA methylation, and survival data were extracted. The basic clinico-pathological features of reserved samples were showed in Supplementary Tab. S1.

### Tumor-infiltrating immune cells analysis

TIMER (https://cistrome.shinyapps.io/timer/) is an integrated web platform for systematic analysis of immune infiltration across various cancer types from TCGA datasets, including 10,897 samples across 32 cancer types [13]. TIMER applies a deconvolution method to speculate the abundance of tumor-infiltrating immune cells (TIICs) according to gene expression profiles [14]. We assessed the correlation between PD-1/PD-Ls expression and methylation with the infiltration levels of TIICs according to infiltrating data obtained from TIMER, including B cells, CD4+ T cells, CD8+ T cells, macrophages, neutrophils, and dendritic cells.

### Co-expression gene and enrichment analysis

To explore genes that shared co-expressed pattern with PD-1/PD-Ls, the online database Linked Omics (http://www.linkedomics.org/login.php) was applied [15], which containing gene expression data with the RSEM normalized count. PD-1/PD-Ls co-expressed genes were analyzed statistically using Pearson’s correlation coefficient. Function module of Linked Omics performs analysis of Gene Ontology (GO) by gene set enrichment analysis (GSEA). The criterion for GO analysis as follows: minimum number of genes was 3, simulations were 500, and analysis method was affinity propagation.

### Statistical analysis

R 3.6.3 and GraphPad Prism 8 were applied as main tools for the statistical analysis and figures exhibition. Most of the data between two groups were analyzed by Student’s t-test. All data are presented in violin plots. Correlation analysis were assessed by Pearson correlation analysis. Survival analysis were conducted in R 3.6.3 by batch analysis methods using self-compiled program, and the groups were divided by median PD-1/PD-Ls gene expression or methylation level. Kaplan-Meier survival plots were generated with survival curves compared by log-rank test. For all analysis, differences were considered statistically significant when P values were less than 0.05.

## Results

### PD⍰1/PD⍰Ls expression levels in LGG

We firstly checked the expression levels of PD⍰1/PD⍰Ls in LGG tissues. As exhibited in Fig. 1A, the expression of these three immune checkpoints in LGG showed obvious clustering. Expression of PD-1/PD-Ls were generally determined in LGG samples. PD-L2 exhibited the highest expression while the expression of PD-1 was lowest among these three immune checkpoints (Fig. 1B). To further identify the correlation between PD⍰1/PD⍰Ls expression, we conducted correlation analysis using Pearson test. The results suggested that PD⍰1/PD⍰Ls had distinct co-expression patterns. PD-1 expression was positively associated with PD-L1 (Fig. 1C) and PD-L2 expression (Fig. 1D), the expression levels between PD-L1 and PD-L2 were also significantly correlated (Fig. 1E). These results uncovered that PD-1, PD-L1, and PD-L2 might collaborate on specific molecular and biological functions in LGG.

**Figure 1.**
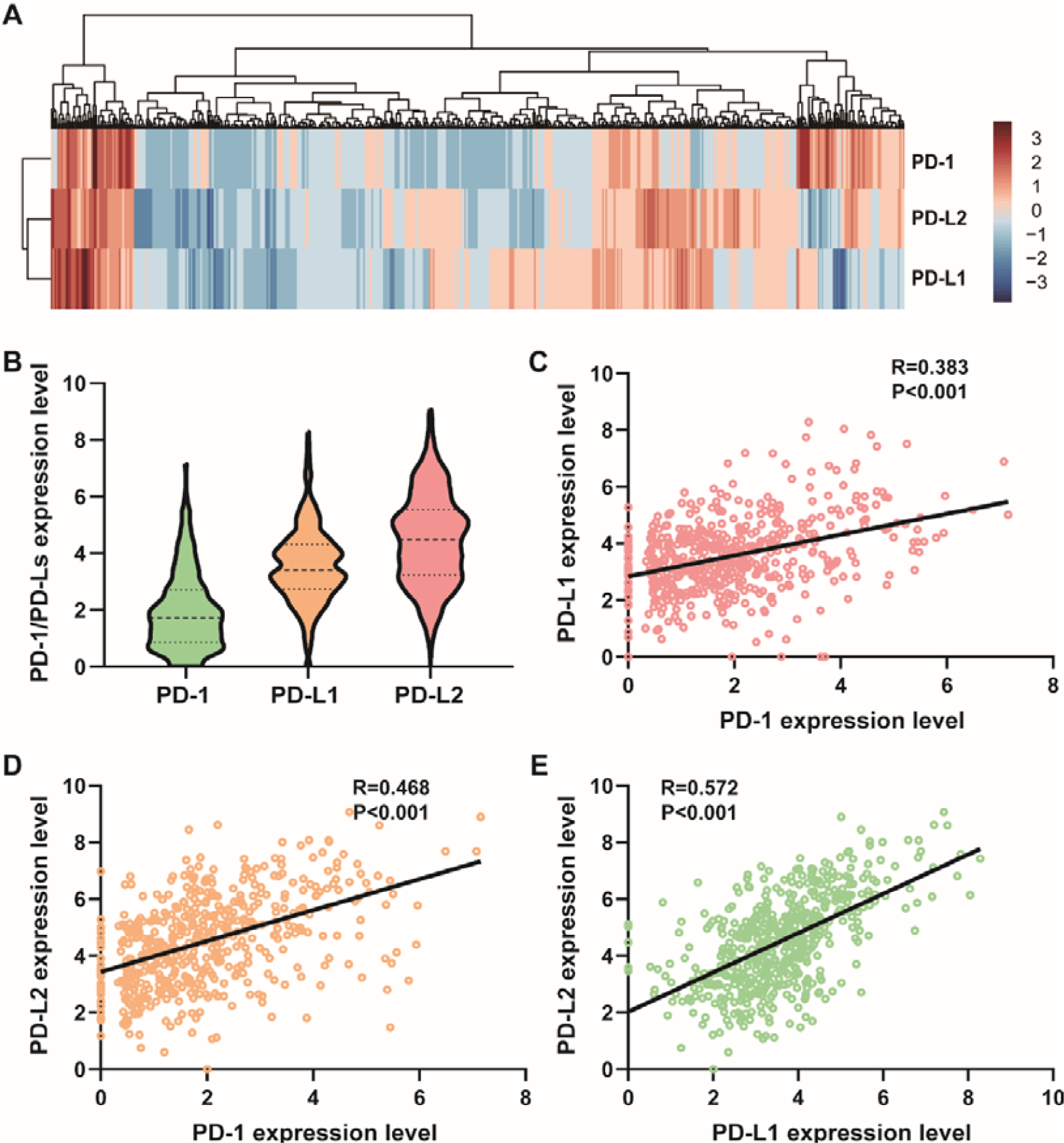
Expression of PD-1/PD-Ls in LGG. (A) A clustering heat map of gene expression of PD-1/PD-Ls in LGG; (B) Expression of PD-1/PD-Ls were generally determined; (C) PD-1 expression was positively correlated with PD-L1 expression; (D) PD-1 expression was positively correlated with PD-L2 expression; (E) PD-L1 expression was positively correlated with PD-L2 expression.

### Clinical significance of PD⍰1/PD⍰Ls expression

Due to notable heterogeneity of molecular nature across different LGG subtypes, PD-1/PD-Ls expression levels were evaluated according to the histological type, WHO grade system, as well as IDH mutation status. The results showed that astrocytoma exhibited the highest PD-1/PD-Ls expression when compared with oligoastrocytoma and oligodendroglioma (Figs. 2A-2C). Moreover, when WHO grade system was applied as a sub-classifier, we found that in grade 3 LGG showed significantly upregulated PD-1/PD-Ls expression (Figs. 2D-2F). Furthermore, IDH mutant-type showed universally lower expression of PD-1/PD-Ls than that of IDH-wt LGG (Figs. 2G-2I). These findings suggested that PD-1/PD-Ls related immune response were obviously various, which may further reflected different biological pattern across LGG subtypes.

**Figure 2.**
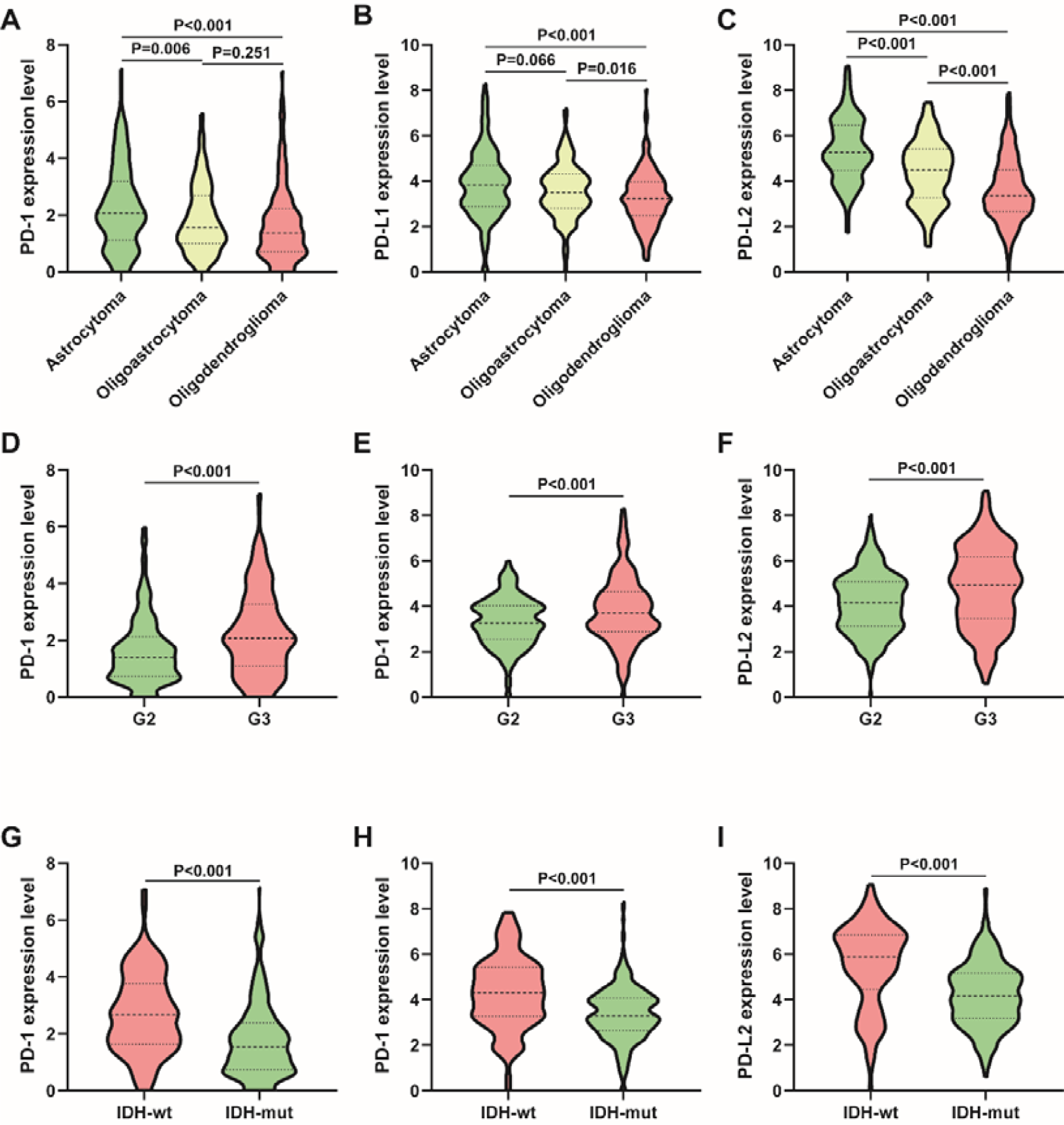
Expression of PD-1/PD-Ls in different LGG subtypes. (A) PD-1, (B) PD-L1, and (C) PD-L2 were highly enriched in astrocytoma subtype; (D) PD-1, (E) PD-L1, and (F) PD-L2 were highly enriched in grade 2 LGG; (G) PD-1, (H) PD-L1, and (I) PD-L2 were highly enriched in IDH wild type LGG.

To acquire the novel insight into the influence of on survival, we checked the prognostic values of PD-1/PDLs expression in LGG. As shown in Fig. 3, patients who expressed higher PD-1 in tumor tissues exhibited a significantly shorter overall survival (OS) than the counterparts (Fig. 3A). Besides, overexpression of PD-L1 (Fig. 3B) and PD-L2 (Fig. 3C) also predicted poor OS in LGG patients. Moreover, we combined these three immune checkpoints expression to evaluate prognosis of LGG patient. The result showed patients with PD-1/PD-Ls high expression had significantly poor OS than other cohorts, and the prognosis of patients with PD-1/PD-Ls low expression was best (Fig. 3D). Overall, these findings suggested that PD-1/PD-Ls were both negative prognostic indicators in LGG.

**Figure 3.**
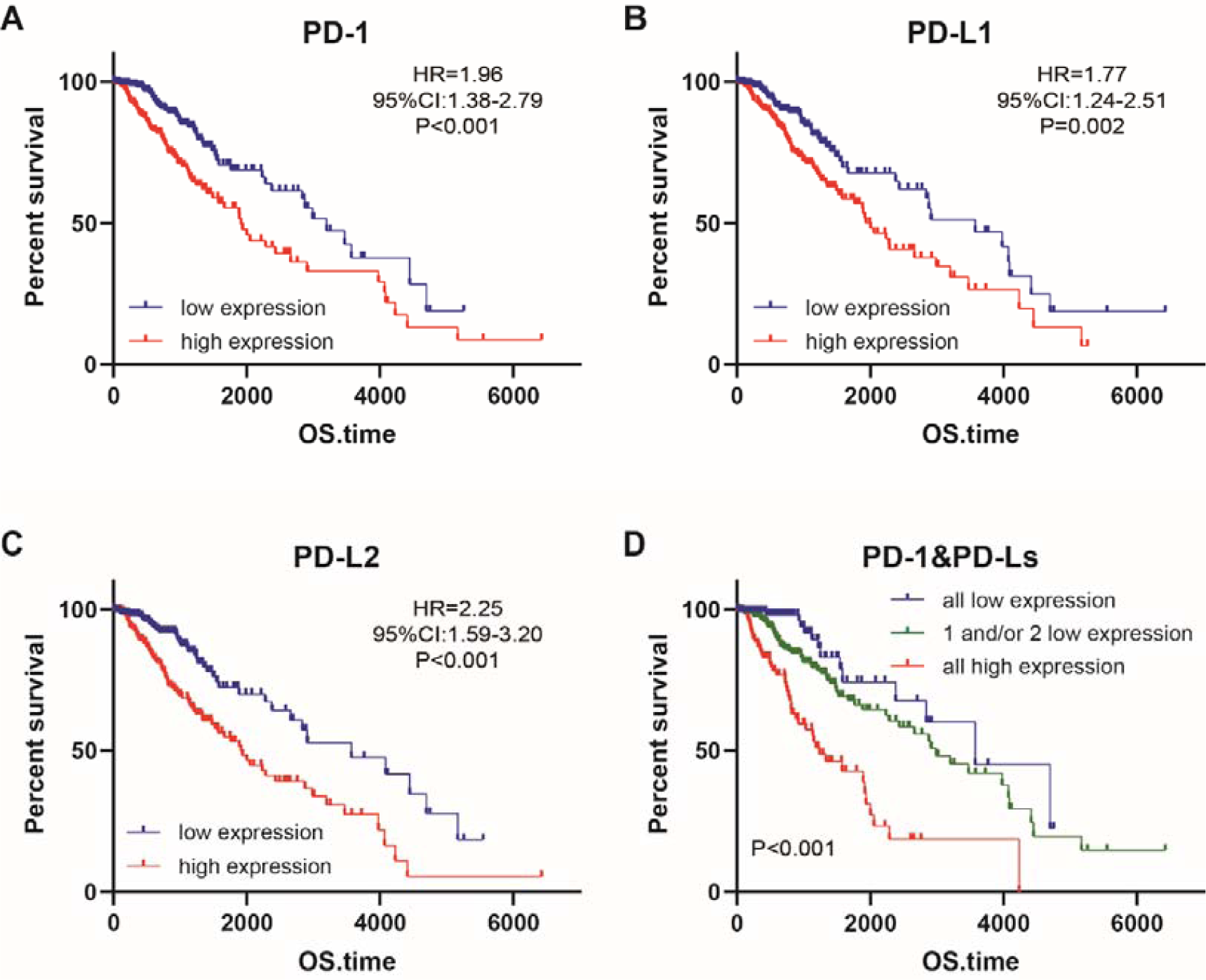
Survival analysis for PD-1/PD-Ls expression. Kaplan–Meier survival analysis showed that high expression of (A) PD-1, (B) PD-L1, and (C) PD-L2 were associated with significantly worse prognosis in LGG patients; (D) Combined PD-1, PD-L1, and PD-L2 expression defined various prognosis in LGG patients.

### Regulatory factors for PD⍰1/PD⍰Ls expression

In view of the fact that PD-1/PD-Ls had various expression patterns in LGG and significant prognostic values were observed, we next try to the regulatory factors that responsible for dys-regulation of PD-1/PD-Ls at gene level according to available data. DNA copy number variations (CNV) are most common genetic alterations that affect tumorigenesis of cancers *via* mediating tumor-related gene expression [16-18]. However, the expression levels of PD-1 and PD-L1 showed no significant difference across different CNV status (Figs. S1A, S1B), while different, CNV status significantly influence PD-L2 expression. Copy gain was associated with notably upregulated PD-L2 levels compared with the copy-neutral (diploid) and copy-loss (shallow deletion and deep deletion) samples (Fig. S1C).

DNA methylation is other common regulatory factor at gene level, which causes low gene expression. We performed DNA methylation clustered based on the expression of PD-1/PD-Ls (Figs. 4A-4C), which showed that most CpG sites had the obviously negative correlation with PD-1/PD-Ls expression. Next, we calculated the mean levels of PD-1/PD-Ls methylations and conducted Pearson correlation analysis to confirm the associations between DNA methylation and mRNA expression. The results exhibited that PD-1 methylation level was negatively associated with PD-1 expression (Fig. 4D). Besides, expression levels of PD-L1 and PD-L2 were regulated by methylation as well (Figs. 4E, 4F). Taken together, DNA methylation was a crucial factor for dysregulated PD-1/PD-Ls expression, which might be used as an indicator for immune checkpoints expression prediction.

**Figure 4.**
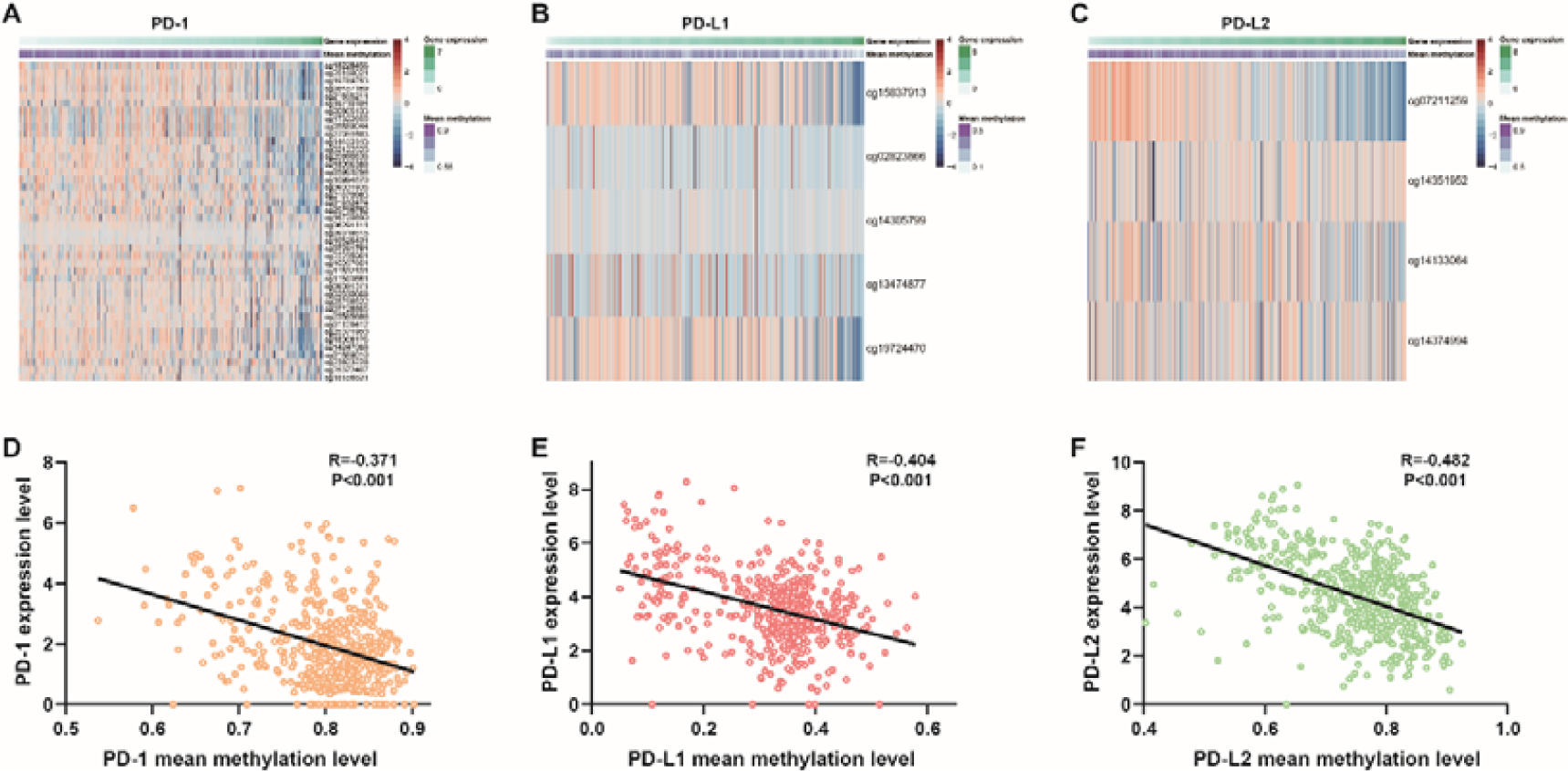
DNA methylation of PD-1/PD-Ls and correlation with expression. The expression clustered methylation of (A) PD-1, (B) PD-L1, and (C) PD-L2; (D) Methylation levels of (D) PD-1, (E) PD-L1, and (F) PD-L2 were negatively with their mRNA expression.

### Clinical significance of PD-1/PD-Ls methylation

Considering the DNA methylation was a crucial factor in regulating PD-1/PD-Ls expression, we further analyzed the methylation levels according to different clinical subtypes. Contrary to gene expression, astrocytoma exhibited the lowest PD-1/PD-Ls methylation levels when compared with oligoastrocytoma and oligodendroglioma (Figs. 5A-5C). Besides, we found that in grade 3 LGG showed significantly hypo-methylation levels than grade 2 (Figs. 5D-5F). Moreover, IDH mutant-type showed commonly hyper-methylation of PD-1/PD-Ls than that of IDH-wt LGG (Figs. 5G-5I). These results suggested that PD-1/PD-Ls methylation levels were potential markers for different LGG subtypes.

**Figure 5.**
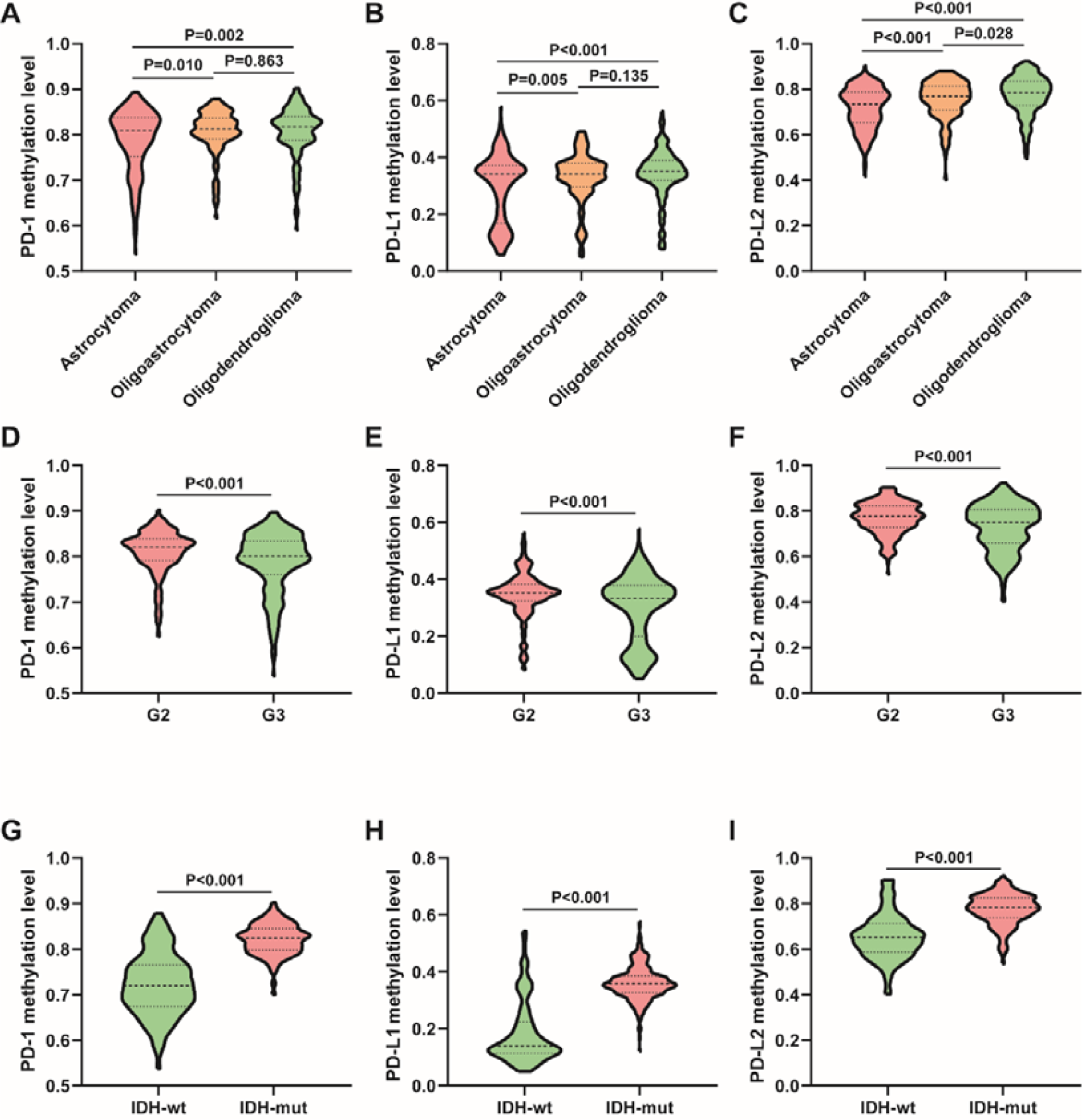
Methylation of PD-1/PD-Ls in different LGG subtypes. (A) PD-1, (B) PD-L1, and (C) PD-L2 were hypo-methylated in astrocytoma subtype; (D) PD-1, (E) PD-L1, and (F) PD-L2 were hypo-methylated in grade 2 LGG; (G) PD-1, (H) PD-L1, and (I) PD-L2 were hypo-methylated in IDH wild type LGG.

DNA methylation has been identified as potential prognostic markers in tumors [19]. Thus, we also checked the prognostic values of PD-1/PDLs methylation in LGG. We first evaluated the prognostic impacts of single CpG site in LGG, the results were exhibited in Tab. 1, and most CpG sites were positive prognostic indicators in LGG. When it came to mean methylation levels, as shown in Fig. 6, patients who with low PD-1 methylation in tumor tissues showed a significantly worse OS than the counterparts (Fig. 6A). Besides, hyper-methylation of PD-L1 (Fig. 6B) and PD-L2 (Fig. 6C) also predicted unfavorable OS in LGG patients. When we combined PD-1/PD-Ls methylations to assess prognosis of LGG patient, the result showed patients with PD-1/PD-Ls hypo-methylations had notably poor OS than other cohorts (Fig. 6D). To sum up, PD-1/PD-Ls methylations might be more effective prognostic indicators than mRNA expressions in LGG.

**Figure 6.**
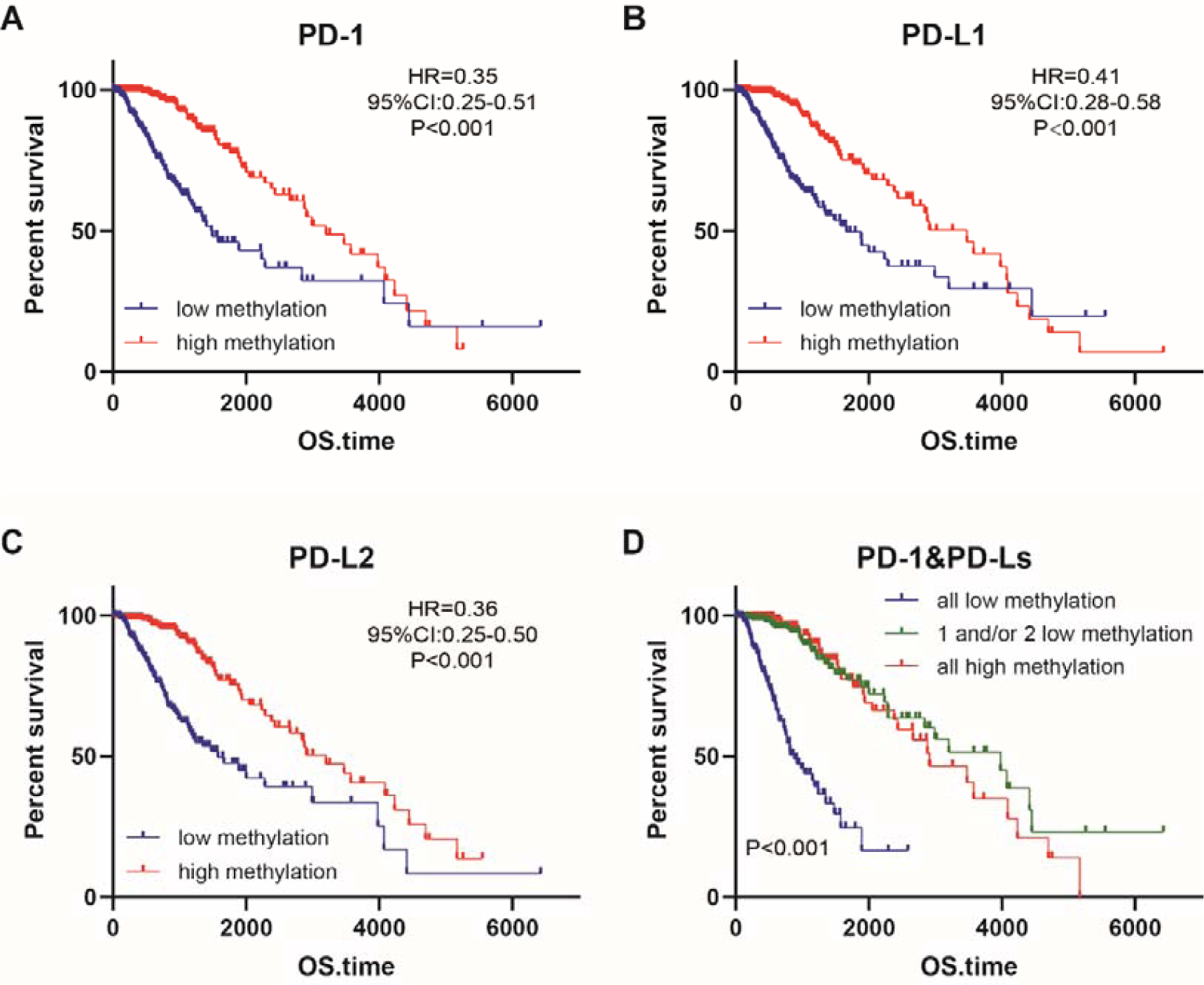
Survival analysis for PD-1/PD-Ls methylation. Kaplan-Meier survival analysis showed that hyper-methylation of (A) PD-1, (B) PD-L1, and (C) PD-L2 were associated with significantly worse prognosis in LGG patients; (D) Combined PD-1, PD-L1, and PD-L2 methylation defined various prognosis in LGG patients.

**Table 1.** Prognostic values of specific CpG sites in PD-1/PD-Ls in LGG.

| **Genes** | **CpG sites** | **HR** | **Low 95%CI** | **Up 95%CI** | **P value** |
| --- | --- | --- | --- | --- | --- |
| PD-1 | cg18228456 | 0.69 | 0.49 | 0.98 | **0.038** |
| PD-1 | cg25150021 | 0.54 | 0.38 | 0.77 | **<0.001** |
| PD-1 | cg19789753 | 0.48 | 0.34 | 0.69 | **<0.001** |
| PD-1 | cg08457169 | 0.59 | 0.41 | 0.84 | **0.002** |
| PD-1 | cg21855211 | 0.52 | 0.37 | 0.74 | **<0.001** |
| PD-1 | cg19710184 | 0.61 | 0.43 | 0.86 | **0.005** |
| PD-1 | cg20805133 | 0.36 | 0.25 | 0.51 | **<0.001** |
| PD-1 | cg17322655 | 0.44 | 0.31 | 0.64 | **<0.001** |
| PD-1 | cg03889044 | 0.51 | 0.35 | 0.72 | **<0.001** |
| PD-1 | cg27051683 | 0.46 | 0.32 | 0.65 | **<0.001** |
| PD-1 | cg14453145 | 0.41 | 0.29 | 0.58 | **<0.001** |
| PD-1 | cg02122525 | 0.39 | 0.27 | 0.56 | **<0.001** |
| PD-1 | cg25890838 | 0.66 | 0.47 | 0.94 | **0.021** |
| PD-1 | cg18096388 | 0.42 | 0.30 | 0.61 | **<0.001** |
| PD-1 | cg03903296 | 0.64 | 0.45 | 0.91 | **0.012** |
| PD-1 | cg10994870 | 0.67 | 0.47 | 0.96 | **0.022** |
| PD-1 | cg09031938 | 0.73 | 0.51 | 1.04 | 0.073 |
| PD-1 | cg21670983 | 0.67 | 0.47 | 0.95 | **0.023** |
| PD-1 | cg01632474 | 0.55 | 0.39 | 0.79 | **0.001** |
| PD-1 | cg25798782 | 0.41 | 0.29 | 0.59 | **<0.001** |
| PD-1 | cg16720890 | 0.62 | 0.43 | 0.88 | **0.007** |
| PD-1 | cg06291111 | 0.61 | 0.43 | 0.87 | **0.006** |
| PD-1 | cg09319815 | 0.89 | 0.63 | 1.26 | 0.498 |
| PD-1 | cg10526431 | 0.75 | 0.53 | 1.07 | 0.111 |
| PD-1 | cg07281781 | 0.77 | 0.55 | 1.10 | 0.152 |
| PD-1 | cg22235901 | 0.81 | 0.57 | 1.16 | 0.232 |
| PD-1 | cg10057601 | 0.60 | 0.42 | 0.86 | **0.003** |
| PD-1 | cg11532131 | 0.51 | 0.36 | 0.73 | **<0.001** |
| PD-1 | cg11503661 | 0.65 | 0.46 | 0.93 | **0.015** |
| PD-1 | cg09391371 | 0.52 | 0.36 | 0.73 | **<0.001** |
| PD-1 | cg02530668 | 0.58 | 0.40 | 0.82 | **0.002** |
| PD-1 | cg03756522 | 0.59 | 0.42 | 0.84 | **0.003** |
| PD-1 | cg07728865 | 0.68 | 0.48 | 0.96 | **0.029** |
| PD-1 | cg23585686 | 0.47 | 0.33 | 0.66 | **<0.001** |
| PD-1 | cg01128412 | 0.60 | 0.42 | 0.85 | **0.004** |
| PD-1 | cg25371950 | 0.44 | 0.31 | 0.62 | **<0.001** |
| PD-1 | cg18308176 | 0.45 | 0.32 | 0.65 | **<0.001** |
| PD-1 | cg14247008 | 0.54 | 0.38 | 0.77 | **0.001** |
| PD-1 | cg01889010 | 0.65 | 0.46 | 0.92 | **0.015** |
| PD-1 | cg23623228 | 0.71 | 0.50 | 1.02 | 0.055 |
| PD-1 | cg25372407 | 0.54 | 0.38 | 0.77 | **0.001** |
| PD-1 | cg18156831 | 0.66 | 0.47 | 0.95 | **0.021** |
| PD-L1 | cg15837913 | 0.31 | 0.22 | 0.45 | **<0.001** |
| PD-L1 | cg02823866 | 0.65 | 0.45 | 0.92 | **0.014** |
| PD-L1 | cg14305799 | 1.08 | 0.76 | 1.54 | 0.669 |
| PD-L1 | cg13474877 | 0.40 | 0.28 | 0.57 | **<0.001** |
| PD-L1 | cg19724470 | 0.38 | 0.27 | 0.54 | **<0.001** |
| PD-L2 | cg07211259 | 0.38 | 0.27 | 0.55 | **<0.001** |
| PD-L2 | cg14351952 | 0.73 | 0.51 | 1.04 | 0.076 |
| PD-L2 | cg14133064 | 0.54 | 0.38 | 0.77 | **<0.001** |
| PD-L2 | cg14374994 | 0.70 | 0.49 | 1.00 | **0.045** |

### Prognostic values of PD-1/PD-Ls in patients with various therapeutic strategies

Radical resection is the preferred therapeutic strategy for LGG, but additional radiotherapy and targeted therapy are important complementary therapeutic strategies [20]. Given the promising prognostic values of PD-1/PD-Ls expression and methylation in LGG, we next try to check the prognostic effects of PD-1/PD-Ls in LGG patients with radiotherapy, without radiotherapy, with targeted therapy, and without targeted therapy. As shown in Tab. 2, methylation levels was the more effective indicators in predicting prognosis of patients receiving different therapeutic strategies than expression levels, which could serve as prognostic indicators in patients with radiotherapy, without radiotherapy, with targeted therapy, and without targeted therapy. However, PD-1/PD-Ls expression also had encouraging prognostic values in specific subgroups. Concretely, high expression of PD-1 was notably associated with poor OS in LGG patients without radiotherapy, with targeted therapy, and without targeted therapy; High expression of PD-L1 were prominently associated with poor OS in LGG patients without radiotherapy and with targeted therapy; Besides, high expression of PD-L2 were significantly associated with poor OS in LGG patients with radiotherapy, with targeted therapy, and without targeted therapy. Taken together, these findings suggested the promising roles of PD-1/PD-Ls as potential prognostic indicators in LGG patients with manifold regimens of therapies.

**Table 2.** Prognostic values of PD-1/PD-Ls expression and methylation in LGG patients with various therapeutic strategies.

| **Indicators** | **Therapeutic strategies** | **HR** | **Low 95%CI** | **Up 95%CI** | **P value** |
| --- | --- | --- | --- | --- | --- |
| PD-1<br>expression | With radiotherapy | 1.36 | 0.91 | 2.05 | 0.134 |
|  | Without radiotherapy | 4.59 | 2.07 | 10.18 | **<0.001** |
|  | With targeted therapy | 1.68 | 1.07 | 2.64 | **0.021** |
|  | Without targeted therapy | 2.09 | 1.11 | 3.95 | **0.025** |
| PD-1<br>methylation | With radiotherapy | 0.39 | 0.26 | 0.59 | **<0.001** |
|  | Without radiotherapy | 0.27 | 0.12 | 0.61 | **0.001** |
|  | With targeted therapy | 0.36 | 0.23 | 0.58 | **<0.001** |
|  | Without targeted therapy | 0.28 | 0.15 | 0.53 | **<0.001** |
| PD-L1<br>expression | With radiotherapy | 1.41 | 0.94 | 2.12 | 0.099 |
|  | Without radiotherapy | 2.34 | 1.07 | 5.12 | **0.041** |
|  | With targeted therapy | 1.65 | 1.05 | 2.58 | **0.029** |
|  | Without targeted therapy | 1.78 | 0.94 | 3.36 | 0.081 |
| PD-L1<br>methylation | With radiotherapy | 0.41 | 0.27 | 0.62 | **<0.001** |
|  | Without radiotherapy | 0.28 | 0.13 | 0.62 | **0.004** |
|  | With targeted therapy | 0.32 | 0.20 | 0.51 | **<0.001** |
|  | Without targeted therapy | 0.36 | 0.19 | 0.68 | **0.002** |
| PD-L2<br>expression | With radiotherapy | 2.55 | 1.70 | 3.85 | **<0.001** |
|  | Without radiotherapy | 1.92 | 0.88 | 4.21 | 0.111 |
|  | With targeted therapy | 2.19 | 1.16 | 4.13 | **0.017** |
|  | Without targeted therapy | 2.31 | 1.48 | 3.62 | **<0.001** |
| PD-L2<br>methylation | With radiotherapy | 0.37 | 0.24 | 0.57 | **<0.001** |
|  | Without radiotherapy | 0.32 | 0.15 | 0.70 | **0.007** |
|  | With targeted therapy | 0.33 | 0.21 | 0.52 | **<0.001** |
|  | Without targeted therapy | 0.32 | 0.17 | 0.63 | **<0.001** |

**Table 3.**
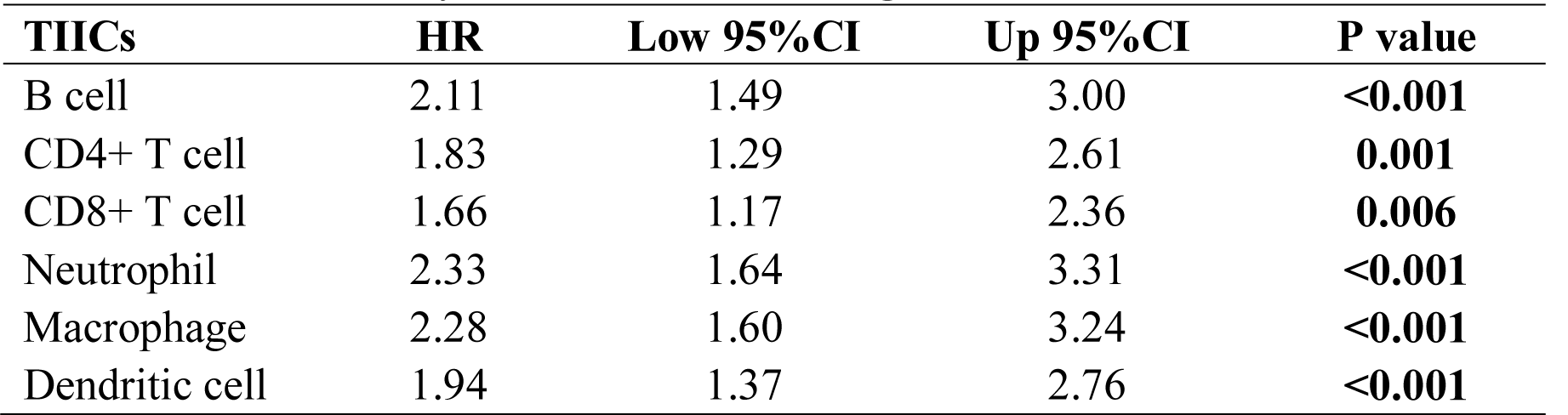
Survival analysis of TIICs infiltrating levels in LGG.

### PD-1/PD-Ls expression as well as methylation and immune infiltration

The survival times of patients in multiple tumors is affected by the quantity and activity status of TIICs [21-23]. As first, we determined the prognostic values of different immune cells infiltration, including B cells, CD4+ T cells, CD8+ T cells, macrophages, neutrophils, and dendritic cells, the results identified that infiltration of these six TIICs all possessed excellent biomarker potential for assessing prognosis (Tab. 3). Next, we explored the relationship between PD-1/PD-Ls expression as well as methylation and the infiltrating immune cells in LGG tissues. The heatmap showed the general correlation between PD-1/PD-Ls expression as well as methylation and infiltration levels of six TIICs (Fig. 7A). Specifically, PD-1, PD-L1, and PD-L2 expression were positively associated with six immune cells infiltrations (Fig. 7B), while PD-1, PD-L1, and PD-L2 were negatively associated with those TIICs infiltrations (Fig. 7B). Therefore, these results further confirmed that PD-1/PD-Ls were specifically correlated with immune infiltrating cells in LGG, suggesting that PD-1/PD-Ls functioned as critical roles in immune escape in the tumor microenvironment.

**Figure 7.**
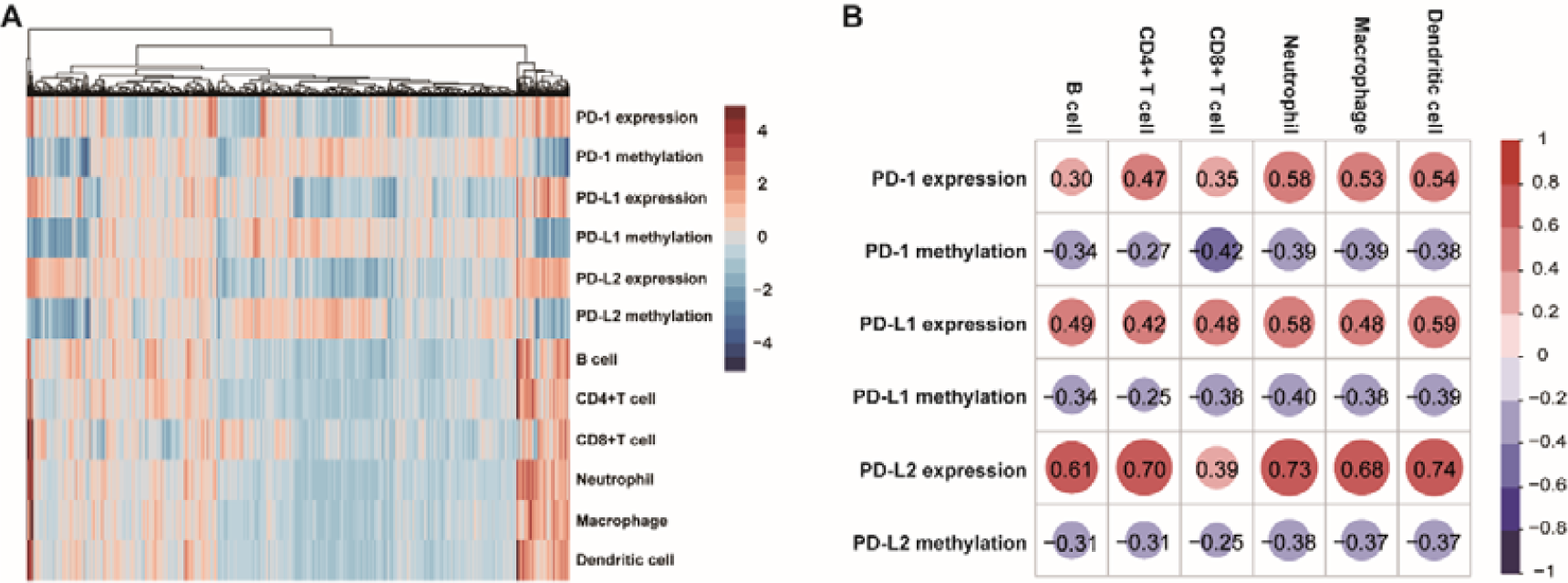
Correlation analysis of PD-1/PD-Ls expression as well as methylation and infiltration levels of immune cells in LGG. (A) Correlation analysis between PD-1/PD-Ls expression as well as methylation and infiltration levels of six TIICs was summarized in the heatmap; (B) PD-1/PD-Ls expression positively correlated with infiltration levels of TIICs, while PD-1/PD-Ls methylation negatively correlated with infiltration levels of TIICs.

### PD-1/PD-Ls associated biological process

To explore the biological features of LGG with different PD-1/PD-Ls expression, we screened the genes that strongly correlated with PD-1/PD-Ls expression, respectively (Figs. 8A-8C). To obtain an exact result, notably related genes were submitted for GO analysis. The results showed that PD-1/PD-Ls co-expressed genes positively regulated immune response related biological processes, such as cellular defense response, adaptive immune response, cellular defense response, and so on (Figs. 8D-8F), while those correlated genes were more related to negatively regulate physiological biological processes, including glutamate receptor signaling pathway, mitochondrial gene expression, etc. (Figs. 8D-8F). Overall, these results revealed that PD-1, as well as PD-L1, PD-L2 were induced as immune inhibitors in the tumor microenvironment where inflammatory and immune response were relatively active, suggesting PD-1/PD-Ls had similar molecular functions in LGG compared with other common solid tumors.

**Figure 8.**
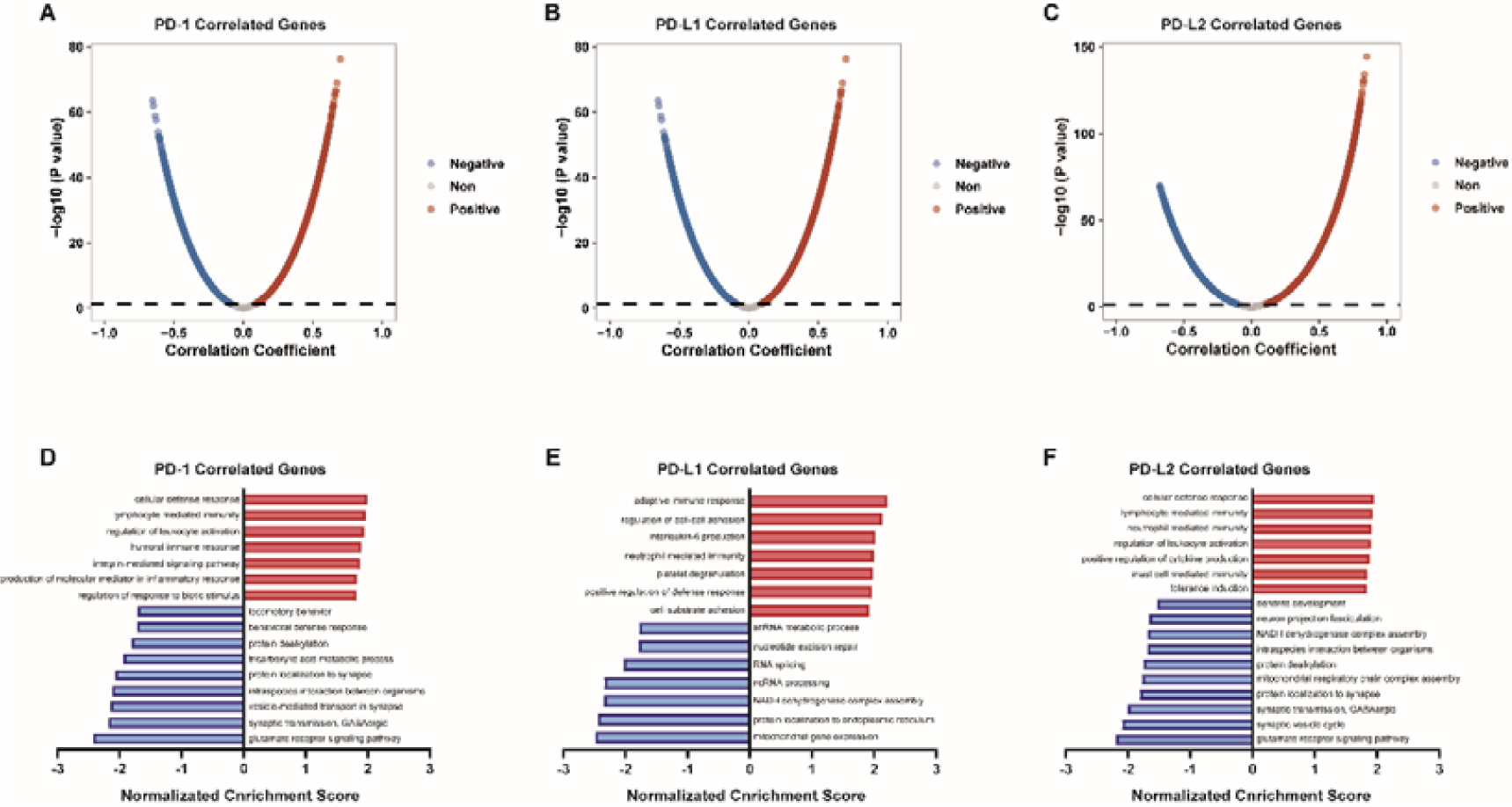
GO analysis of PD-1/PD-Ls co-expressed genes in LGG. The global (A) PD-1, (B) PD-L1, and (C) PD-L2 highly correlated genes identified by Pearson test in LGG. (A) Significantly enriched BP annotations of (D) PD-1, (E) PD-L1, and (F) PD-L2 in LGG.

## Discussion

LGG is the main subtype of gliomas, which characterized with lower aggressiveness and well differentiation than GBM counterpart. A growing number of studies focus on the interplay between glioma and immunity, but most research attach great attentions to the GBM [24-26]. LGG seems to be ignored from the field of cancer immunotherapy due to its better prognosis than GBM. Several studies have uncovered that GBM tend to express higher PD-L1 than LGG [27, 28]. However, whether PD-1/PD-Ls play key roles in LGG and whether patients could benefit from immunotherapy should be explored to further improve curative effect and prognosis.

Immune checkpoints play crucial roles in tumor immune escape. PD-1/PD-Ls axis is the most important immune checkpoints in cancer immunity, but whether PD-1/PD-Ls have notable influence on LGG biology is largely unknown. We have noticed two studies presented by Prof. Jiang’s group reported that PD-L1 and PD-L2 were correlated with WHO grade system and poor prognosis in gliomas and GBM, as well as PD-L2 also had promising prognostic value in LGG [9, 10]. In this research, we systematically evaluated PD-1/PD-Ls expression and their prognostic values in LGG. PD-1/PD-Ls had distinct co-expression patterns and were upregulated in astrocytoma subtype and higher grade LGG. Moreover, expression levels of PD-1/PD-Ls were companied by IDH mutation, indicating that IDH wild-type LGG exhibited more tumor-derived immune response than IDH mutant LGG. Additionally, PD-1/PD-Ls high expression were both correlated with poor prognosis in LGG patients, which could be served as promising prognostic indicators. All results suggested that PD-1/PD-Ls expression was associated with more aggressive biological process in LGG.

We further explored the regulatory factors responsible for the dys-regulated PD-1/PD-Ls expression. In this research, DNA methylation levels were found to be negatively correlated with PD-1/PD-Ls expression, and PD-L2 expression also was regulated by CNVs. In Berghoff *et al*.’s report, PD-L1 expression was negatively mediated by methylation level of CpG site cg15837913 [29], revealing DNA methylation was a significant regulatory factor for PD-L1 expression in LGG. However, RNA-sequencing could not reflect the cellular resources of expression data, which might not explain the precise mechanism of PD-1/PD-Ls expression regulation. Moreover, we also noticed that PD-1/PD-Ls methylations were more effective indicators for LGG subtypes and prognosis. According to previous studies, PD-L1 methylation mediated PD-L1 expression and functioned as a promising prognostic marker in melanoma [30]. Besides, PD-L1 promoter methylation was also associated with negative PD-L1 expression, and the development of advanced gastric cancer [31]. Therapeutic subtype analysis revealed that the methylation of PD-1/PD-Ls had more broad-spectrum prognostic values in LGG patients receiving various therapeutic strategies. Collectively, PD-1/PD-Ls methylation could be innovative biomarkers for assessing LGG patients’ prognosis in addition to PD-1/PD-Ls expression.

Immune checkpoints commonly play significant roles in triggering the cancer immunity escape by mediating the interaction between tumor cells and TIICs, so their expressions were commonly positively correlated with TIICs abundance [32, 33]. Deng *et al*. established an immune-related prognostic signature and found that LGG patients with high-risk had higher levels of infiltrating B cells, CD4+ T cells, CD8+ T cells, macrophages, neutrophils, and dendritic cells [34], indicating TIICs infiltration was an unfavorable prognostic biomarker in LGG. Hao *et al*. identified that infiltration of TIICs was associated with poor prognosis in specific types of LGG [35]. In our research, we found that the high infiltration of TIICs predicted worse outcomes in total LGG patients. Besides, PD-1/PD-Ls expression and methylation were both correlated TIICs infiltration. Although PD-1/PD-Ls methylation exhibited more obviously promising prognostic values, their expression had tighter association with TIICs infiltration, suggesting the various roles of expression and methylation in assessing prognosis and immune infiltration, respectively.

We subsequently identified co-expressed genes of PD-1/PD-Ls to summarize their potential biological functions in LGG and the GO enrichment analysis suggested PD-1/PD-Ls co-expressed genes enriched in immune response related biological processes, which conformed to the defined roles of PD-1/PD-Ls in multiple solid cancers. In view of the similar roles of PD-1/PD-Ls and adverse prognostic TIICs in LGG, we speculated that patients who expressed higher PD-1/PD-Ls notably blocked the anti-tumor effect of TIICs. In other words, those patients might benefit more from immunotherapy.

## Conclusion

In this research, we reported the expression and methylation status of PD-1/PD-Ls in different subtypes of LGG. High PD-1/PD-Ls expression and hypo-methylation of PD-1/PD-Ls were associated with poor survival of LGG patients. PD-1/PD-Ls expression were demonstrated to be related with immune cells infiltration. Besides, the PD-1/PDLs correlated gene profiles were screened, the GO enrichment analysis of which focus on immune response related biological process. To sum up, LGG patients with PD-1/PD-Ls high expression, whose prognosis was poorer, might benefit from PD1/PD-Ls checkpoint inhibitors treatment.

## Abbreviation

WHO: the World Health Organization;
LGG: low-grade glioma;
GBM: glioblastoma;
PD-1: programmed death 1;
CD279: cluster of differentiation 279;
PD-L1: programmed death-ligand 1;
PD-Ls: PD-L1 and PD-L2;
TCGA: the Cancer Genome Atlas;
TIICs: tumor-infiltrating immune cells;
GO: gene ontology;
GSEA: gene set enrichment analysis;
OS: overall survival;
CNV: copy number variations;

## Declarations

### Ethics approval and consent to participate

Not applicable.

### Consent for publication

Not applicable.

### Availability of data and material

All data are included in the article.

## Authors’ contributions

CL, HW, and JM conceived the study and participated in the study design, performance, coordination and manuscript writing. JM, YC, RX, XY, and WZ carried out the assays and analysis. CL and HW revised the manuscript. All authors reviewed and approved the final manuscript.

## Acknowledgements

Not applicable.

## Figure legends

**Figure S1.**
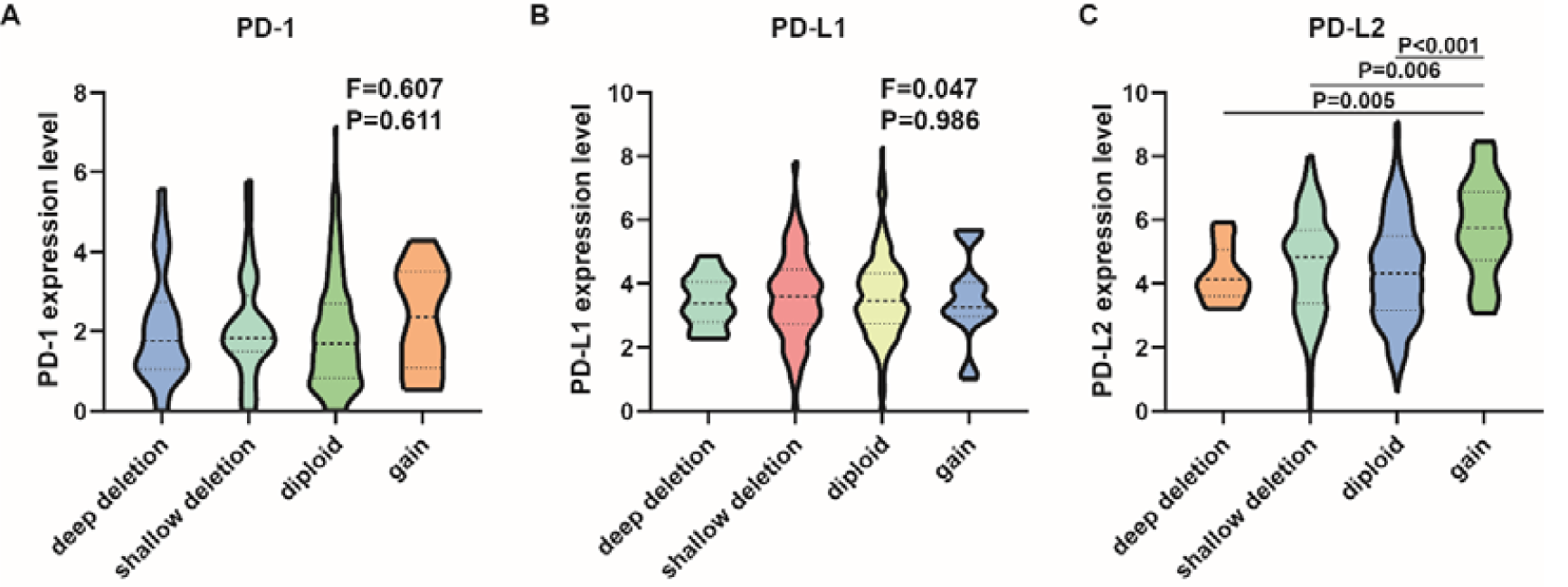
Correlation between the CNVs of PD-1/PD-Ls and expression levels in LGG tissues. CNVs in (A) PD-1, (B) PD-L1 had no association with their expression; (C) Copy gain of PD-L2 was associated with notably upregulated expression level compared with the copy-neutral and copy-loss cases.

**Table S1.**
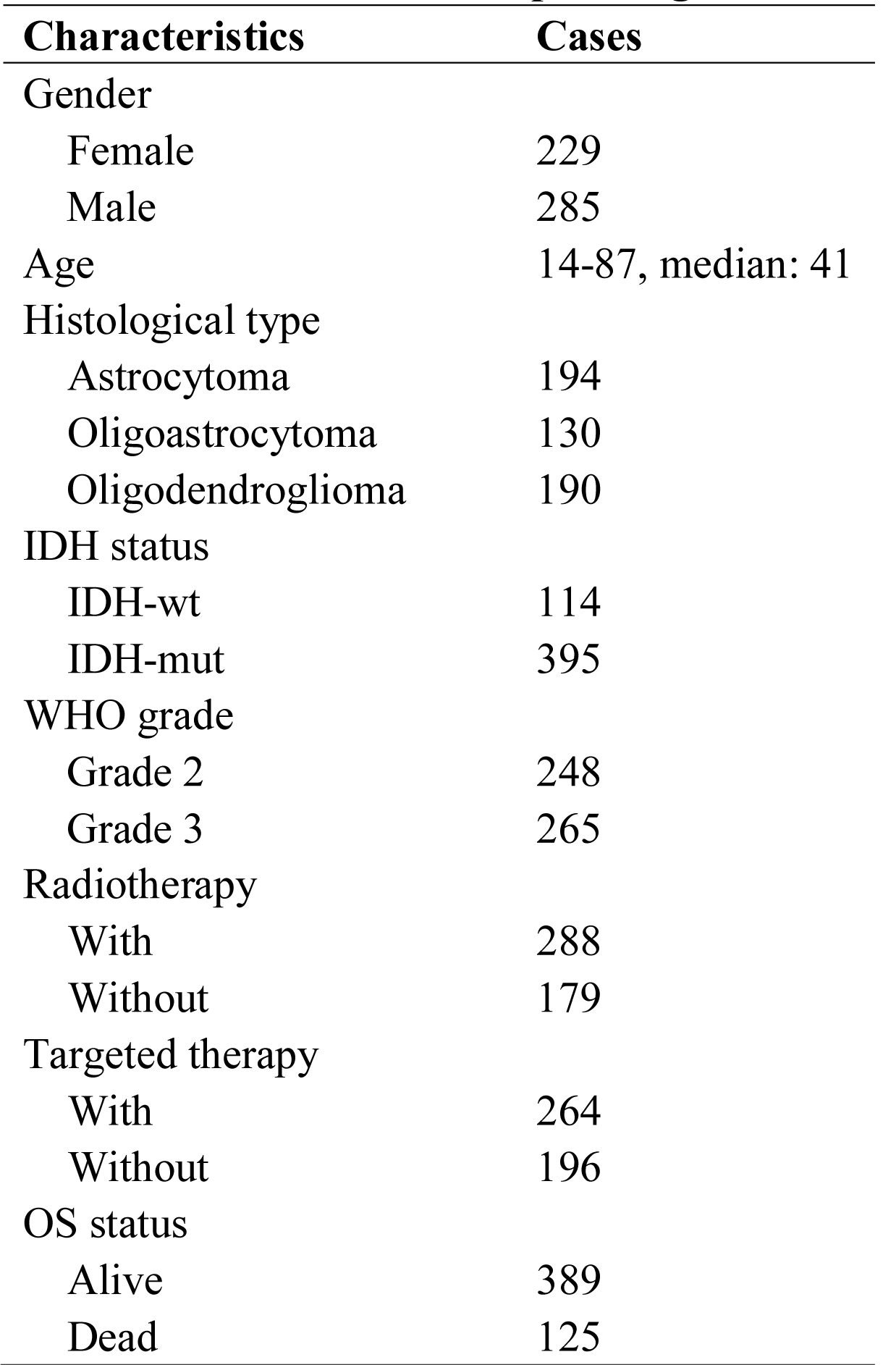
The basic clinico-pathological features of LGG patients.

